# Development of a Versatile System to Facilitate Targeted Knockout/Elimination Using CRISPR/Cas9 for Highly Duplicated Gene Families in *Arabidopsis* Sexual Reproduction

**DOI:** 10.1101/2024.04.22.590670

**Authors:** Hidenori Takeuchi, Shiori Nagahara

**Author notes:** Corresponding author: Hidenori Takeuchi. Present address: Division of Biological Sciences, Graduate School of Science, Kyoto University, Kyoto 606-8502, Japan (Shiori Nagahara).

## Abstract

CRISPR/Cas9-based targeted gene editing is a fundamental technique for studying gene functions in various organisms. In plants, the introduction of a T-DNA construct harboring Cas9 nuclease and single guide RNA (sgRNA) sequences induces sequence-specific DNA double-strand breaks, inducing the loss of gene function. *Arabidopsis thaliana* is a model for CRISPR/Cas9 system development and gene function studies; the introduction of *Cas9* under the egg or zygote promoter and multiple sgRNA modules generates heritable or non-mosaic mutants for multiple targets in the T1 generation of *A. thaliana*. Recent reports reflect use of several CRISPR/Cas9 vectors in generating single– and higher-order mutants; however, the development of a reliable, cost-effective, and high-throughput CRISPR/Cas9 platform is necessary for targeting highly duplicated gene families. In this study, we have developed a simple and user-friendly construction system for the CRISPR/Cas9 vector series with improved gene editing efficiency by simply inserting a single intron into *Cas9*, and effectively demonstrated the simultaneous knockout of multiple genes involved in *A. thaliana* sexual reproduction. An unbiased PCR-mediated mutant identification in the T1 generation revealed that our CRISPR/Cas9 system can support a > 70 kb deletion of > 30 tandemly duplicated synergid-specific genes and simultaneous knockout of five redundant genes essential for double fertilization. We performed a one-shot knockout of seven homologous pollen tube receptor-like kinase genes and identified their specific and overlapping roles in pollen tube growth and guidance. Our system can potentially facilitate further research in experimental plant biology to search for genetically unidentified components using reverse genetic candidate approaches.

## Introduction

The CRISPR/Cas9 system that presents a tool for precise gene mutagenesis has revolutionized genetic research in a wide range of organisms. To generate precise genomic mutations in mammalian cells, this system was first developed based on the discovery of programmable RNA-guided DNA endonucleases involved in bacterial immunity; subsequently, it was applied to various organisms (Jinek et al. 2012, Cong et al. 2013, Mali et al. 2013, Wang and Doudna 2023), including model plants, such as the *Arabidopsis thaliana* (dicot), *Oryza sativa* (monocot), and the liverwort *Marchantia polymorpha*, as well as crop species (Feng et al. 2013, Mao et al. 2013, Brooks et al. 2014, Sugano et al. 2014, Xing et al. 2014, Endo et al. 2015). CRISPR/Cas9 system involves *Agrobacterium*-mediated genetic transfer of a T-DNA construct encoding the Cas9 nuclease and a single guide RNA (sgRNA) into plant cells; the Cas9-sgRNA complex is assembled in plant cells and stochastically induces double-strand breaks in target sequences, which is directed by 20 nucleotides present in the 5’ end of the sgRNA, resulting in a non-homologous end-joining (NHEJ)-mediated short insertions or deletions at the targeted loci. Cas9 expression and DNA cleavage in cells, such as egg, zygote, and meristematic cells, which produce the entire plant body, can lead to heritable mutations (Wang et al. 2015, Tsutsui and Higashiyama 2017). Direct transfer of the Cas9-sgRNA complex into plant cells has been reported to generate CRISPR/Cas9-mediated gene-edited plants (Woo et al. 2015, Toda et al. 2019) although this is still challenging owing to the difficulties in efficient Cas9-sgRNA complex introduction, regeneration, and selection to establish individual plants.

Using *A. thaliana*, CRISPR/Cas systems have been developed in terms of their efficiency and multiplicity. These systems effectively support gene function studies. Moreover, CRISPR/Cas-based technology is effective in inducing relatively large (>10 kb) deletions, chromosomal rearrangements, gene targeting (i.e., gene knock-in), and epigenetic regulation (Ordon et al. 2016, Miki et al. 2018, Wolter and Puchta 2019, Beying et al. 2020, Grützner et al. 2021, Jogam et al. 2022). Previous studies indicate that the efficiency of CRISPR/Cas-based technology significantly depends on the type and expression level of Cas protein as well as the selection of sgRNA targets. Therefore, promoters driving codon-optimized Cas proteins and vector construction systems for the introduction of multiple sgRNAs have been explored and several useful vectors were developed to generate gene-edited plants (Wang et al. 2015, Yan et al. 2015, Tsutsui and Higashiyama 2017, Grützner et al. 2021, Ursache et al. 2021). However, for the reverse genetic approach, a more efficient CRISPR/Cas9 system associated with a simple and user-friendly platform can facilitate generating multiple mutants required.

Duplication of genes involved in intercellular and inter-organism communication is generally common and correlated with certain selective pressures for subfunctionalization (Swanson and Vacquier 2002). In particular, genes involved in sexual reproduction (intercellular male-female communication) potentially impact molecular evolution and occasionally show characteristic gene multiplication. For instance, reproductive tissues express hundreds of genes encoding secreted cysteine-rich peptides that are grouped into several subfamilies (Bircheneder and Dresselhaus 2016, Takeuchi 2021, Okuda et al. 2009, Sprunck et al. 2012, Takeuchi and Higashiyama 2012, Zhong et al. 2019). The functions of a single male gametophyte cell, the pollen tube, are regulated by more than ten receptor-like kinases belonging to several subfamilies (Muschietti and Wengier 2018, Muschietti et al. 1998, Miyazaki et al. 2009, Boisson-Dernier et al. 2009, Liu et al. 2013, Chang et al. 2013, Takeuchi and Higashiyama 2016, Wang et al. 2016, Ge et al. 2017, Zhou et al. 2021). Therefore, a reliable method for the simultaneous knockout of multiple genes would be beneficial. However, the available CRISPR/Cas9 system and platforms, which involve time-consuming and labor-intensive processes, are insufficient to address this problem.

In this study, we have developed a simple vector construction system targeting CRISPR/Cas9-mediated multiple gene knockout and analyzed its utility for highly duplicated gene families associated with *A. thaliana* sexual reproduction. We established an system that involves a set of binary Cas9 vectors with two promoters and three antibiotic selection markers, and a set of subcloning vectors for reliable assembly of up to 12 sgRNA expression units in a binary vector. Using this system, we demonstrated that the insertion of a single intron into the *Cas9* gene significantly improved the efficiency of gene editing, leading to the generation of mutants with more than a 70 kb deletion, knockout of five redundant genes essential for double fertilization, or one-shot knockout of seven homologous pollen tube genes that regulate pollen tube growth and guidance. This study paves the way for the identification of genetically unidentified components using reverse genetic approaches, particularly for gene families consisting of redundant duplicated genes.

## Results and Discussion

### A simple and reliable construction system for introducing multiple sgRNA modules into the CRISPR/Cas9 vector

To search for genetically unidentified components of interest, a user-friendly CRISPR/Cas9 system and a simple, reliable, and stable platform for establishing a collection of candidate mutants are desirable; which include (1) CRISPR/Cas9 constructs that can be easily removed from the plant genome after genome editing, (2) a simple cloning procedure for introducing multiple sgRNA modules into a binary vector series, and (3) a convenient and cost-effective method for identifying Cas9-mediated mutants of multiple genes. First, we inserted the red fluorescent seed selection cassette, *At2S3::mCherry* (Kroj et al. 2003), into the CRISPR/Cas9 vector (pHEE401E, Wang et□al.□2015), which harbored an egg cell-specific promoter for *zCas9* (*Cas9* codon-optimized for *Zea mays*) and *Bsa*I restriction sites for Golden Gate assembly of sgRNA modules (Fig. 1A, designated as “pHEE-R-hyg”). A previous study showed that different colors of fluorescent seed selection cassettes were useful for selecting individuals co-transformed with two CRISPR/Cas9 vectors (Ursache et al. 2021). Considering the complicacy associated with the selection of dual fluorescent seeds in T1 and non-fluorescent seeds in T2 using different filters of the fluorescence stereomicroscope, we used multiple antibiotic selection markers. We generated a Cas9 vector series containing kanamycin-resistance (*npt* II) or bialaphos-resistance (*bar*) genes by replacing the hygromycin-resistance gene (*hpt* II) of pHEE-R-hyg; thus, the pHEE-R-kan and pHEE-R-bar vectors were generated (Supplementary Table S1).

**Fig. 1.**
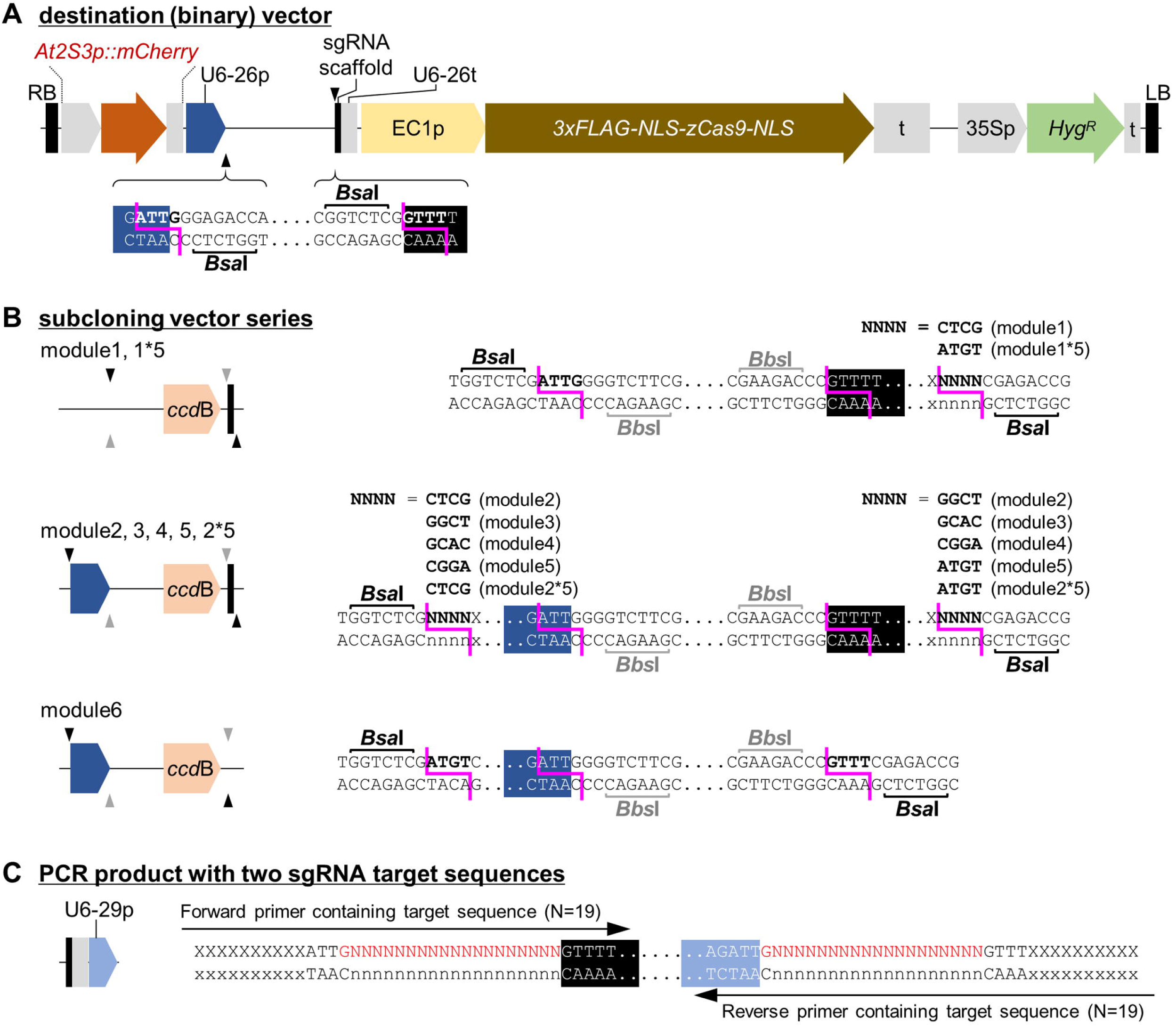
Schematic representation of CRISPR/Cas9 vector construction for multiple sgRNA modules. (A) pHEE401E-based CRISPR/Cas9 binary vector containing the red fluorescent seed selection cassette, *At2S3::mCherry*. RB, right border; LB, left border; ‘p’, promoter; ‘t’, terminator; *Hyg^R^*, hygromycin resistance gene. (B) Subcloning vector series, modules 1, –2, –3, –4, –5, –6, –1*5, and 2*5. Black and gray triangles indicate the position of *Bsa*I and *Bbs*I restriction sites, respectively. (C) The first and second sgRNA target sequences added to both ends of the PCR product for cloning into the subcloning vectors.

Initially, we directly used PCR-amplified gene fragments to clone multiple sgRNA modules into a binary vector through the Golden Gate assembly. However, considering occasional low efficacies in the assembly and sequence error(s) in one of the modules, we prepared a subcloning vector series for each sgRNA module, suitable to be sequentially connected and introduced into the Cas9 vector series (Fig. 1B). Two sgRNA expression units were prepared using a simple one-step PCR, following the previously reported system of pHEE401E (Wang et□al.□2015), and inserted into each subcloning vector by subsequent seamless cloning or Golden Gate assembly (Fig. 1C, Supplementary Method). CRISPR/Cas9 vector harboring 12 sgRNA expression units can be constructed by Golden Gate assembly using subcloning vector series (‘module1’, ‘module2’, ‘module3’, ‘module4’, ‘module5’, and ‘module6’ as shown in Fig. 1B) and one of the binary vectors. In cases of fewer targets, 4 sgRNA expression units (using ‘module1*5’ and ‘module6’) and 6 sgRNA expression units (using ‘module1’, ‘module2*5’, and ‘module6’) are also assembled in the same way. This multiplexed sgRNA assembly system supports easy incorporation of multiple sgRNAs targeting two or more sites per gene or locus and disruption of gene structure through deletion/elimination (several hundred to several kilobase pairs). This strategy facilitates easy and inexpensive screening of the genome-edited lines based on variable band sizes detected through conventional PCR of the target gene followed by agarose gel electrophoresis (as described later).

### High-efficiency gene editing via insertion of a single intron of *UBIQUITIN10* between the promoter and the *Cas9* coding region

*Cas9* expression as well as the design and target loci of sgRNAs significantly influence the gene editing efficiency. Previous studies reported that changing *zCas9* to 13 intron-containing z*Cas9* (*zCas9i*) improves gene editing efficiency (Grützner et al. 2021, Ursache et al. 2021). Similarly, we established a method to improve *Cas9* expression through simpler sequence addition, the insertion of a single intron from *A. thaliana UBIQUITIN10* (*UBQ10*), which increases both mRNA and protein levels (Rose 2004). We inserted the *UBQ10* single intron (304 nt; from –304 to –1 nucleotides from the start codon of an endogenous *UBQ10*) with an additional nuclear localization signal (NLS) sequence at the junction between the promoter and the start codon of *zCas9* (Fig. 2A). Additionally, we created the *A. thaliana RPS5A* promoter version by swapping the egg cell-specific promoter of the original vector (Fig. 2A), considering that both promoters are effective for generating heritable Cas9-mediated mutants (Wang et□al.□2015, Tsutsui and Higashiyama, 2017). We constructed a series of CRISPR/Cas9 vectors in combination with the three different antibiotic selection markers: pHEE-R-hyg9si, pHEE-R-kan9si, pHEE-R-bar9si, pHEE-R-hyg5A, pHEE-R-kan5A, pHEE-R-bar5A, pHEE-R-hyg5A9si, pHEE-R-kan5A9si, and pHEE-R-bar5A9si (Supplementary Table S1).

**Fig. 2.**
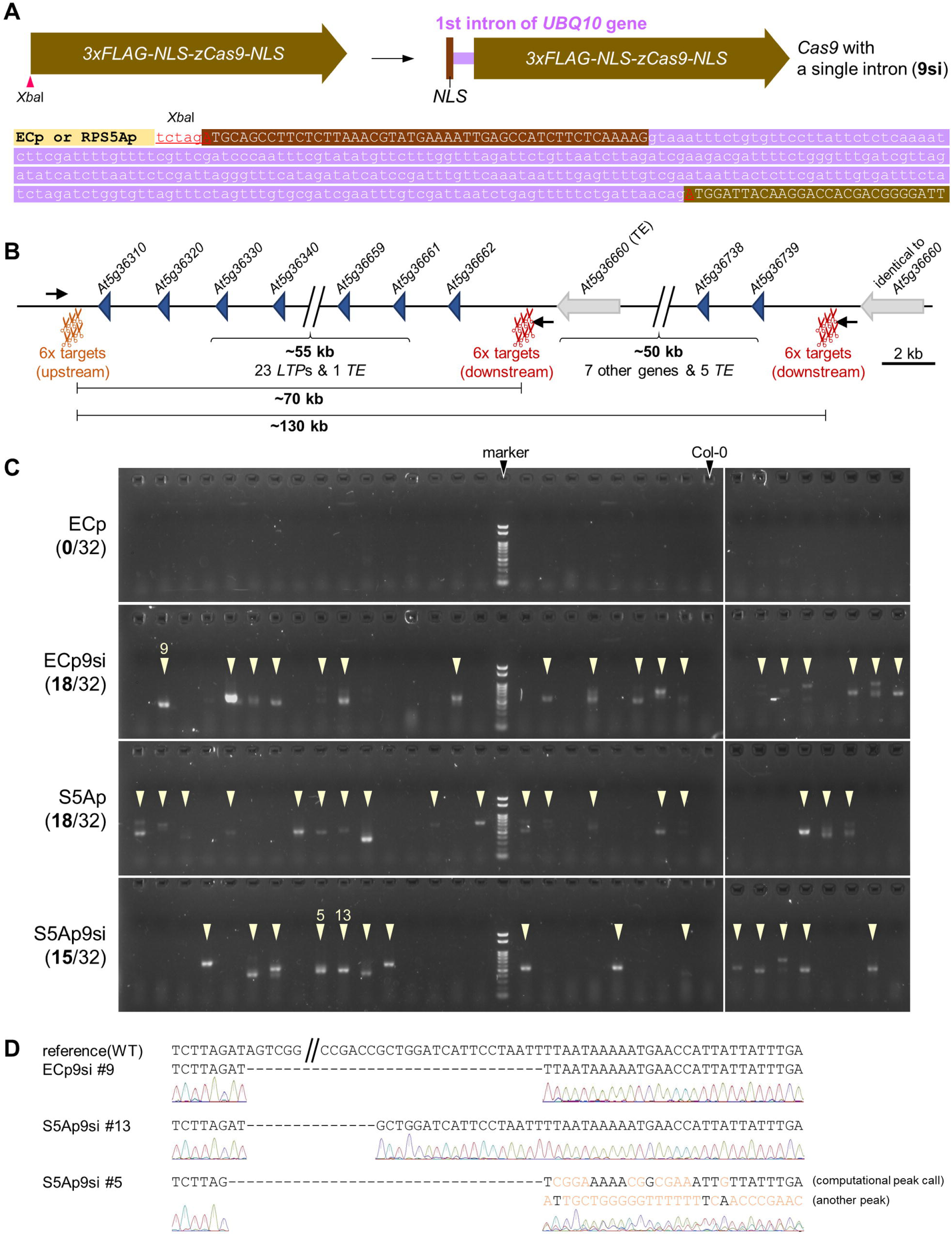
Improvement of genome editing efficiency using *UBQ10* intron with *NLS* sequence inserted between the promoter and the *Cas9* cassette. (A) Introduction of the single intron (‘si’) of *UBQ10* gene using *Xba*I restriction site. (B) Schematic presentation of a tandemly duplicated synergid-specific *LTP* locus. Arrows indicate sites corresponding to locus-specific primers. Note that the region for the downstream 6 sgRNA targets and primer sequences is not unique; a nearly identical duplicate sequence exists ∼50 kb apart. (C) Agarose gel electrophoretic analysis for PCR-based genotyping to detect deletions in the *LTP* locus. Arrowheads indicate the deletion bands regardless of clear or weak signals. (D) Sequencing analysis of PCR products acquired from three lines; 72317, 72299, and 72319 bp deletions (or 131616, 131598, and 131618 bp deletions if sequence eliminations occurred between the upstream and second downstream target sequences) are shown in ECp9si #9, S5Ap9si #13, and S5Ap9si #13, respectively.

To assess the impact of intron insertion and promoter selection on Cas9-induced gene editing efficiency, we targeted a cluster of numerous tandemly duplicated genes encoding lipid transfer protein (LTP)-related or Early Culture Abundant 1 (ECA1) gametogenesis-related cysteine-rich peptides that are predominantly expressed in synergid cells of female gametophytes (Silverstein et al. 2007, Jones-Rhoades et al. 2007, Susaki et al. 2021). Thirty tandemly duplicated synergid-specific *LTP* genes (*At5g36310* to *At5g36662*) spanning approximately 70 kb were considered. To maximize the probability of deletions at the *LTP* locus, six sgRNAs were designed to target the upstream and downstream regions (Fig. 2B). The downstream 6 sgRNAs targeted a nearly identical duplicate located ∼50 kb apart, in the flanking region of 2 related *LTP* genes, including *At5g36738* and *At5g36739*) (Fig. 2B). The 12 sgRNA units were assembled into each of four binary vectors, including pHEE-R-bar, pHEE-R-bar9si, pHEE-R-bar5A, and pHEE-R-bar5A9si, using the subcloning vectors modules 1, 2, 3, 4, 5, and 6. We transformed wild-type *A. thaliana* (Col-0) plants using each construct, selected red fluorescent seeds, and genotyped the T1 seedlings. The expected deletion (∼70 kb or ∼130 kb) was detected based on the appearance of a ∼500–1000 bp PCR product (Fig. 2B). None of the 32 T1 individuals generated using pHEE-R-bar (ECp) showed deletions, whereas deletions were identified in 56% (18/32) of the T1 individuals generated with pHEE-R-bar9si (ECp9si) (Fig. 2C). Moreover, pHEE-R-bar5A (S5Ap) and pHEE-R-bar5A9si (S5Ap9si)-mediated transformation led to the deletions in 56% (18/32) and 46% (15/32) T1 individuals, respectively (Fig. 2C). Most of the PCR bands in ECp9si and S5Ap9si were clear, whereas approximately half of the PCR bands in S5Ap appeared faint (Fig. 2C), suggesting the stochastic deletions in a small fraction of leaves in S5Ap lines, leading to the generation of somatic mosaics (Tsutsui and Higashiyama, 2017). Direct sequencing of the PCR products confirmed deletion of sequence longer than 70 kb (Fig. 2D). These data demonstrate that the insertion of a single intron improves the gene editing efficacy and the combination of multiple sgRNAs enables targeted large chromosomal deletions (more than 70 kb) via a conventional method of transforming wild-type *A. thaliana*.

### One-shot generation of multiple gene knockout mutants: a case study of five duplicated *EGG CELL 1* (*EC1*) genes required for double fertilization

Subsequently, to verify the effectiveness of our system in vector construction and identification of multiple gene knockout mutants, we targeted five *EC1* genes, *EC1.1* (*At1g76750*), *EC1.2* (*At2g21740*), *EC1.3* (*At2g21750*), *EC1.4* (*At4g39340*), and *EC1.5* (*At5g64720*), which encode small cysteine-rich peptides secreted by the egg cell and are redundantly involved in sperm membrane fusion with the egg and central cells in the double fertilization process (Sprunck et al. 2012, Cyprys et al. 2019, Wang et al. 2024). Twelve sgRNAs were designed to induce small targeted deletions in five *EC1* genes located at four genomic loci (Fig. 3A). We transformed Col-0 plants with each of the four constructs (ECp, ECp9si, S5Ap, and S5Ap9si) containing the 12 sgRNA units; genotyping PCR revealed occurrence of the expected gene deletions in the T1 lines generated using ECp9si, S5Ap, and S5Ap9si (Fig. 3B, Supplementary Fig. 1 and 2). We further focused on T1 lines with a ∼1.5 kb deletion in the *EC1.2-1.3* locus, and selected candidate quintuple *ec1* mutants (Fig. 3B, Supplementary Fig. 1 and 2). Although direct sequencing of the PCR products can not always validate Cas9-induced mutant sequences owing to complex biallelic mutations, no wild-type sequences of *EC1* were detected in these candidates, indicating potential loss of functional *EC1* genes in these candidates.

**Fig. 3.**
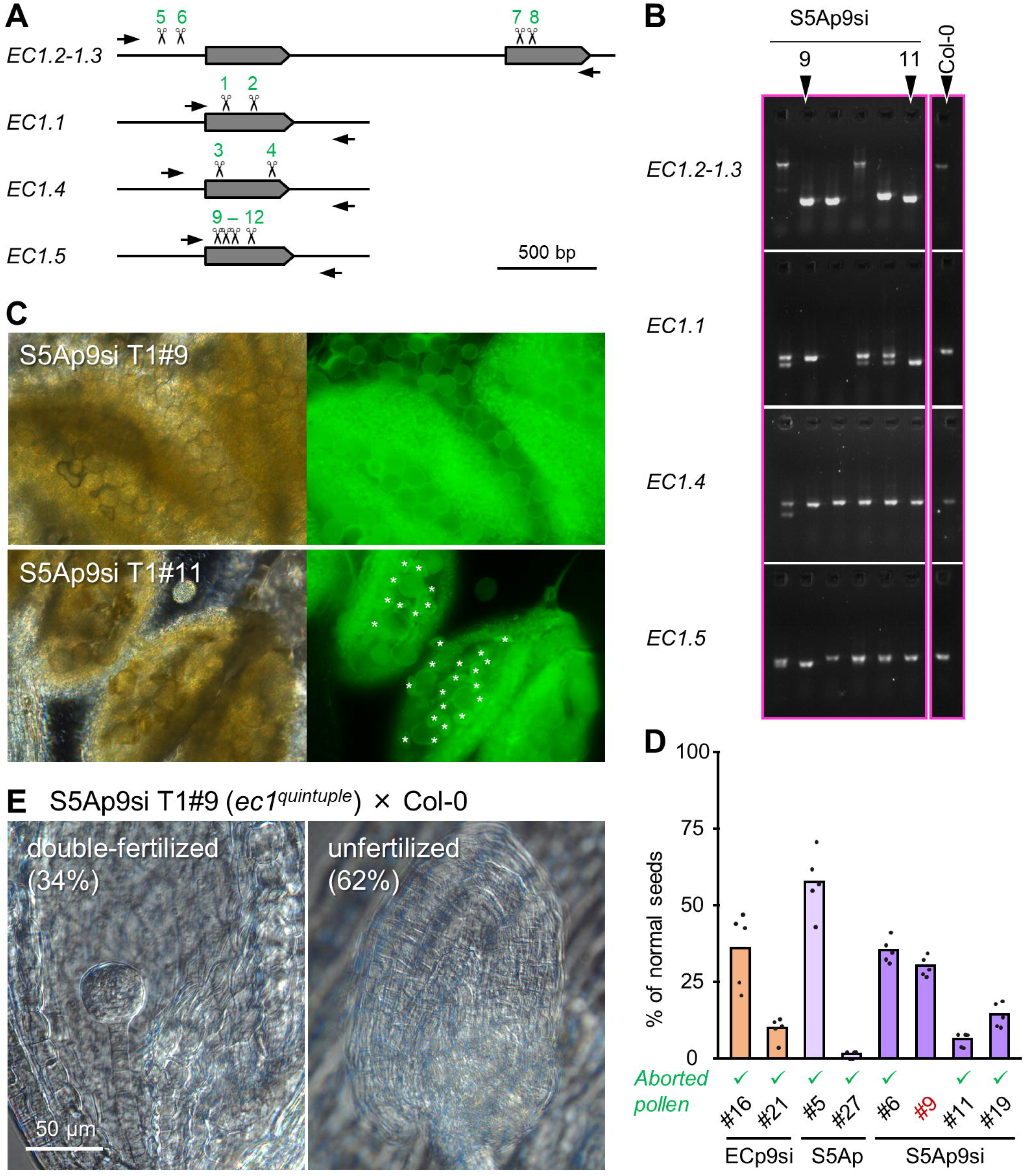
One-shot knockout of five redundant *EC1* genes. (A) Schematic presentation of four loci for five *EC1* genes. Arrows indicate the positions of gene-specific primer sequences. Scissors indicate the target sites of sgRNAs. Numbers (1-12) correspond to the order of sgRNA modules in the CRISPR/Cas9 binary vector. (B) Portions of agarose gel electrophoresis images depicting the results of PCR-based genotyping of the four *EC1* loci. For all gel images, see Supplementary Fig. 1 and 2. (C) Microscopic images of pollen grains in the anther by light field (left) and epifluorescence (right). Anthers in S5Ap9si #9 contained normal pollen grains, whereas S5Ap9si #11 contained collapsed aborted pollen grains (asterisks in epifluorescence image). (D) The percentage of normal developing seeds in auto-pollinated siliques of T1 plants. Dots represent the individual values for analyzed siliques (*n* = 5). Checkmarks indicate T1 lines with aborted pollen grains. (E) Successful (double-fertilized) and failed (unfertilized) embryo and endosperm developments in cleared ovules three days after pollination. Percentages of double-fertilized and unfertilized ovules detected using 239 ovules obtained from 6 pistils are indicated in the panels.

*Agrobacterium*-mediated transformation often induces chromosomal translocations or rearrangements leading to lethality in both male and female gametophytes without inducing somatic abnormalities (Ray et al. 2017, Nacry et al. 1998, Clark and Krysan 2010). As transformants with abnormal gametophyte development show reduced seed fertility, the verification of normal development in gametophytes should be conducted before analyzing the detailed phenotypes associated with the disrupted reproductive genes. Hence, we examined whether pollen (male gametophytes) development was normal using pollen autofluorescence following our simple screening scheme (Fig. 3C). One (S5Ap9si #9) of the eight selected candidate lines exhibited normal pollen grains in the anthers, while the other 7 lines had abnormally collapsed pollen grains (Fig. 3C and D). A percentage of normal seeds was 30% in the siliques of S5Ap9si #9 plants (Fig. 3D), which is the expected fertility in quintuple *ec1* mutants (Sprunck et al. 2012, Wang et al. 2024). Lower percentages (<30%) of normal seeds were observed in the siliques of some of the other lines (Fig. 3D), suggesting that gametophyte lethality in those other lines affected fertility. We assessed embryo and endosperm development after pollen tube arrival using the S5Ap9si #9 line as females. Expectedly, 62% (*n* = 239) of the ovules were unfertilized in the quintuple *ec1* mutant (S5Ap9si #9), which was indicated by the absence of developing embryos or endosperm (Fig. 3E). These results confirm the effectiveness of our system, especially the Cas9 vectors with *UBQ10* single intron, for simultaneous multiple gene knockouts, even in the T1 generation established using wild-type Col-0. As Cas9-mediated mutant generation via *Agrobacterium*-mediated transformation may induce chromosome rearrangements more frequently, mutant candidates should be cautiously and carefully screened, especially when focusing on reproductive phenomena.

### One-shot generation of multiple gene knockout mutants: an investigation of seven homologous genes of *POLLEN RECEPTOR-LIKE KINASE* (*PRK*) family required for pollen tube growth/guidance

We applied our system to the search for genetically unidentified components; hence, we targeted a group of pollen-specific receptor-like kinases, the PRK family (Chang et al. 2013, Takeuchi and Higashiyama 2016). In *A. thaliana*, seven of the eight *PRK* genes (*PRK1*, *2*, *3*, *4*, *5*, *6*, and *8*) are specifically and abundantly expressed in the pollen tube, and therefore, they are considered to play an essential role in pollen tube function. Previously, we generated several multiple *prk* mutants by crossing T-DNA insertion mutants based on their phylogenetic relationships, resulting in various higher-order mutants, including quadruple mutants (*prk1 prk2 prk4 prk5* and *prk1 prk3 prk6 prk8*), and observed a significant reduction in seed fertility, pollen tube growth, and guidance (Takeuchi and Higashiyama 2016). Here, we generated *de novo* septuple *prk* mutants using the CRISPR/Cas9 system to elucidate the collective function of seven pollen-expressing *PRK* family genes in *A. thaliana* sexual reproduction. To maximize the probability of deletions in each gene, three sgRNAs for *PRK1*, *PRK2*, *PRK4*, and *PRK5*, and four sgRNAs for *PRK3*, *PRK6*, and *PRK8* were designed (Fig. 4A). For the simultaneous introduction of 24 sgRNAs, a set of 12 sgRNAs targeting *PRK1*, *PRK2*, *PRK4*, or *PRK5* were intgegrated in pHEE-R-kan (ECp) or pHEE-R-kan9si (ECp9si), and another set of 12 sgRNAs targeting *PRK3*, *PRK6*, or *PRK8* were intgegrated in pHEE-R-hyg (ECp) or pHEE-R-hyg9si (ECp9si). Col-0 plants were transformed with two pairs of constructs (ECp or ECp9si) by co-dipping and T1 transformants were selected on medium containing kanamycin and hygromycin. Eight from the ECp-transformant and five from the ECp9si-transformants were recovered and grown in soil; genomic PCR revealed deletions in some *PRK* genes in two lines (ECp #6 and ECp9si #2) (Fig. 4B). Interestingly, although the other lines exhibited no obvious deletion in target *PRK* genes, the ECp #6 line showed deletions in five targets (*PRK2*, *PRK4*, *PRK5*, *PRK3*, and *PRK6*) and insertion of *PRK1*, and the ECp9si #2 line exhibited deletions in five targets (*PRK1*, *PRK4*, *PRK3*, *PRK6*, and *PRK8*). This suggested that the presumably high Cas9 expression in these transformants induced simultaneous DNA cleavage at most of the target loci. By direct sequencing of the PCR products, we confirmed the absence of wild-type *PRK* sequences in ECp #6 and ECp9si #2 lines and detected deletions of several hundred bp and 1 bp insertions/deletions, which were later validated using the T2 generation (Supplementary Fig. 3A). For instance, in the ECp #6 line, one allele of the *PRK6* gene had a 426-bp deletion by excision with the first and third sgRNAs, and a 1-bp deletion at the fourth sgRNA (Supplementary Fig. 3B). Surprisingly, the *PRK1* gene exhibited a 404-bp jumping insertion of a *PRK3* fragment excised by the first and third sgRNAs (Supplementary Fig. 3B), which suggested that Cas9-sgRNA-directed DNA cleavage events occur simultaneously in the zygote.

**Fig. 4.**
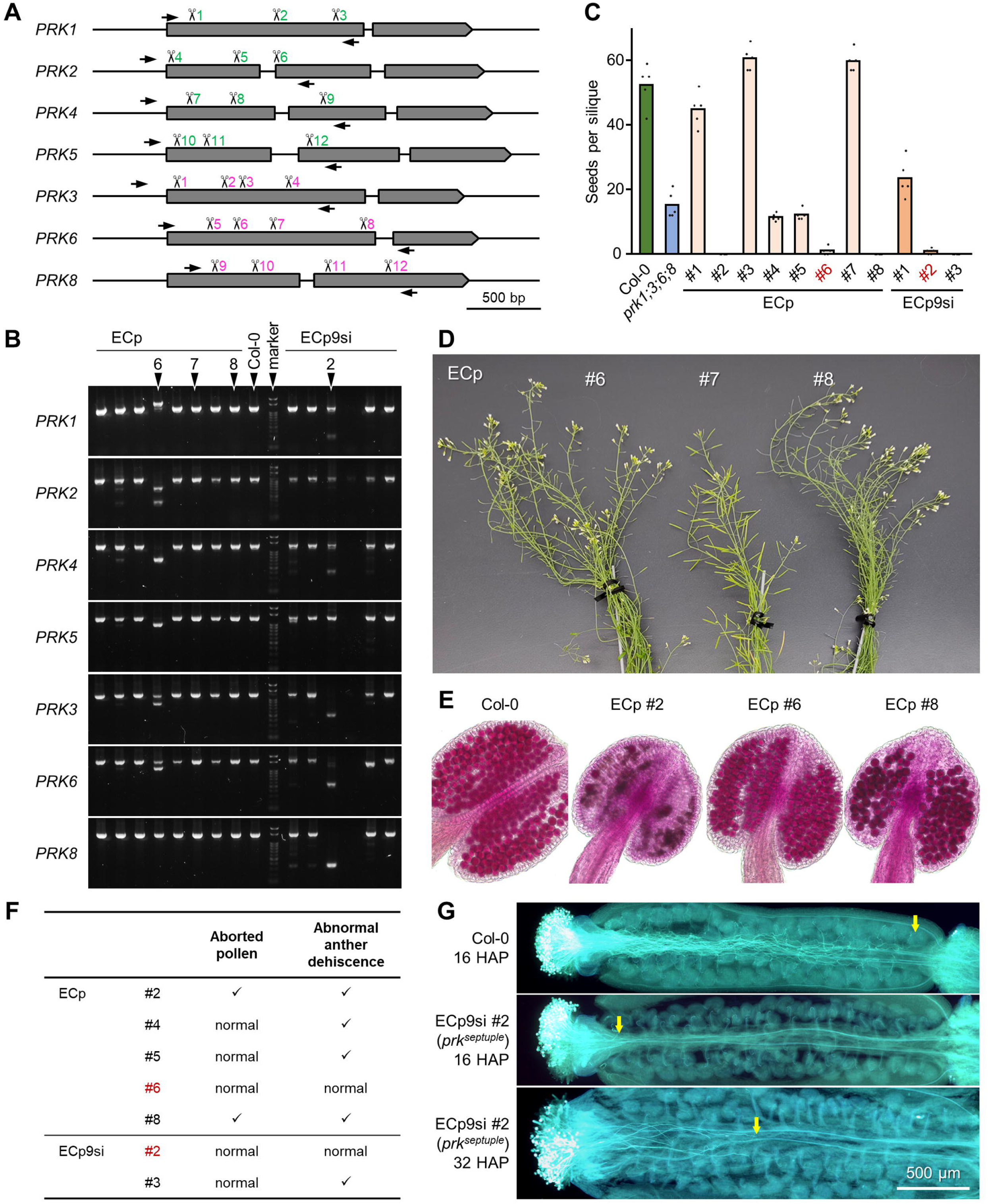
One-shot knockout of seven *PRK* genes. (A) Schematic presentation of seven pollen-expressed *PRK* genes. Arrows indicate the positions of gene-specific primer sequences. Scissors indicate target sites of sgRNAs. Numbers (1-12) marked in green and magenta correspond to the order of sgRNA modules in two independent CRISPR/Cas9 binary vectors. (B) Agarose gel electrophoresis images depicting the results of PCR-based genotyping of the seven *PRK* loci. (C) The number of seeds in auto-pollinated siliques of T1 plants. Dots represent the individual values for analyzed siliques (*n* = 5). (D) Flowers and siliques of three T1 plants (ECp #6, 7, and 8). Line #7 showed almost full seed set, while #6 and 8 showed nearly complete infertility. (E) Pollen viability staining showing normal (red) (Col-0 and ECp #6) and collapsed (dark red) (ECp #2 and #8) pollens. (F) Summarized observation of aborted pollen grains and abnormal anther dehiscence in T1 lines with low seed fertility. (G) Aniline blue staining of pistils 16 and 32 hours after pollination (HAP) with Col-0 or the *de novo* septuple *prk* mutant (ECp9si #2). In each pistil, arrows indicate the tips of a single pollen tube of maximum length.

We counted the number of seeds in the siliques of the eight ECp and three ECp9si T1 lines (Fig. 4C). In this study, wild-type Col-0 and a previously generated quadruple *prk* mutant (*prk1 prk3 prk6 prk8*) produced approximately 50 and 15 seeds per silique, respectively. Despite normal vegetative growth, five T1 lines (ECp #2, #6, and #8; ECp9si #2 and #3) showed a further reduced or no seed fertility (Fig. 4D). However, the ECp #2 and #8 lines had aborted pollen, indicating an undesired abnormality caused by the chromosomal rearrangements as mentioned previously (Fig. 4E). Additionally, the ECp9si #3 line exhibited abnormal anther dehiscence (Fig. 4F), which was also observed in some transgenic lines, leading to reduced fertility. In contrast, normal pollen development and anther dehiscence were evident in ECp #6 and ECp9si #2 lines (Fig. 4E and 4F). The results of genotyping, sequencing, and phenotypic analyses in the T1 generation indicated the successful generation of two (2/13) *de novo* septuple *prk* mutant lines through the CRISPR/Cas9 system.

We analyzed the pollen tube growth in the pistil of these septuple *prk* mutants (Fig. 4G). Sixteen hours after pollination, the wild-type pollen tubes reached the bottom of the transmitting tract of the ovary and targeted most ovules, whereas the longest pollen tube of the septuple *prk* mutant was still around the style. Although several septuple *prk* mutant pollen tubes were detected in the transmitting tract at 32 hours after pollination, they target only < 1% ovules (data from 5 and 7 pistils pollinated by ECp #6 and ECp9si #2 pollen, respectively). In terms of seed fertility, pollen tube growth, and guidance, the phenotype of the septuple *prk* mutant was more severe than that of the quadruple *prk1 prk3 prk6 prk8* mutant (Takeuchi and Higashiyama 2016). However, we could not detect the pollen tube burst phenotype observed in *anx1/2* or *bups1* mutants for another type of receptor-like kinases (Miyazaki et al. 2009, Boisson-Dernier et al. 2009, Ge et al. 2017, Zhou et al. 2021). These outcomes suggest that the seven PRK receptors play specific and overlapping roles in successive steps of pollen tube growth and guidance. Our data demonstrate that our high-throughput CRISPR/Cas9 vector-based strategy is useful in reverse genetic approaches for identifying highly duplicated gene families with uncharacterized functions.

## Conclusions

In conclusion, we modified the egg cell promoter-based pHEE401E vector (Wang et□al.□2015) and developed a CRISPR/Cas9 system effective for the targeted disruption of multiple genes (Fig. 1). This improved system includes a ∼300 nt *UBQ10* single intron added to the original *Cas9* without introns, which significantly enhanced the efficiency of gene editing (Fig. 2). The simple insertion of a relatively short sequence between the promoter and the coding sequence is an easy process, hence, this strategy is considered useful for various CRISPR/Cas-based technologies and heterologous gene expression systems.

After introducing the T-DNA constructs via floral dip transformation of *A. thaliana*, we identified putative mutants through an unbiased PCR-mediated method using all T1 lines without phenotypic pre-selection. Simultaneous expression of multiple sgRNAs, presumably accompanied by high Cas9 levels, induced multiplex gene disruption in certain individuals (Fig. 3B, Fig. 4B, Supplementary Fig. 1 and 2). Therefore, the implementation of another sgRNA inducing a phenotypically selectable mutation, such as trichomeless *gl1* mutation (Miki et al. 2018), in the CRISPR/Cas9 vector can potentially reduce the effort required for candidate screening of the T1 seedlings. In addition to generating the previously reported *ec1* quintuple mutant as a demonstration experiment (Fig. 3), we generated *prk* septuple mutants with a distinct and specific phenotype in pollen tube growth and guidance (Fig. 4). Our data illustrate strategies for rapid generation of multigene knockouts to analyze sexual reproduction-related genes avoiding the pitfalls. The established CRISPR/Cas9 platform provides a reliable, cost-effective, and high-throughput method to facilitate the targeted knockout of highly duplicated genes belonging to a subfamily and eliminate more than 100 kb of genomic loci, thus encouraging further research in plant genetics through the easy generation of a collection of mutants.

## Materials and Methods

### Plasmid construction

All plasmids were generated using standard molecular cloning techniques, including restriction digestion of the backbone vectors followed by ligation with T4 DNA ligase (Nippon Gene, Japan) or the Gibson assembly reaction with Gibson Assembly Master Mix (New England Biolabs Japan, E2611) (Supplementary Table S1). Plasmids pHEE401E and pCBC-DT1T2 were gifted by Qi-Jun Chen (Wang et□al.□2015, Addgene plasmid #71287; Xing et al. 2014, Addgene plasmid #50590). All binary and subcloning vector series generated in this study will be deposited in Addgene (https://www.addgene.org/).

For constructing the binary vector series, the red fluorescent seed selection cassette *At2S3::mCherry* (Kroj et al. 2003) was amplified and inserted into pHEE401E using the *Pme*I and *Hin*d III restriction sites, resulting in pHEE-R-hyg (HTv993). The kanamycin and bialaphos resistance gene cassettes were amplified using vectors derived from pMDC100 and pMDC123 (Curtis and Grossniklaus 2003) and inserted into pHEE-R-hyg by replacing the hygromycin resistance gene cassette at *Mfe*I and *Sac*II restriction sites, thus generating pHEE-R-kan (HTv1562) and pHEE-R-bar (HTv1563), respectively. The *RPS5A* promoter sequence was inserted into pHEE-R-hyg, pHEE-R-kan, and pHEE-R-bar by replacing the egg cell promoter with *Spe*I and *Xba*I restriction sites, resulting in pHEE-R-hyg5A (HTv1564), pHEE-R-kan5A (HTv1565), and pHEE-R-bar5A (HTv1566), respectively. The *UBQ10* intron with the *NLS* sequence was prepared using *A. thaliana* genomic DNA through PCR and inserted into each vector using the *Xba*I restriction site, resulting in pHEE-R-hyg9si (HTv1758), pHEE-R-kan9si (HTv1759), pHEE-R-bar9si (HTv1760), pHEE-R-hyg5A9si (HTv1761), pHEE-R-kan5A9si (HTv1762), and pHEE-R-bar5A9si (HTv1763).

For constructing the subcloning vector series, pUC19-based pT7-Blue (Novagen) was used as the vector backbone. Fragments of the U6-26 promoter and the sgRNA scaffold sequence were amplified from pCBC-DT1T2 and assembled with the backbone sequence in which the *Bsa*I site in the original sequence was mutated. Additional sequences, including *Bbs*I restriction sites for sgRNA target sequence introduction and *Bsa*I plus 4 nt sequences for Golden Gate assembly, were introduced using primers to generate the first version of our subcloning vectors. The *ccdB* cassette was added to the first series using *Bbs*I sites, resulting in the subcloning vector series used: pT7-module1, –2, –3, –4, –5, –6, –1*5, and 2*5 (HTv1567-1574).

Binary vectors containing multiple sgRNA expression units were constructed (Supplementary Method, Supplementary Table S1). The online tool CRISPRdirect (Naito et□al.□2015, http://crispr.dbcls.jp/) was used to select specific target sequences (a single target in *A. thaliana* genome suitable for ‘12mer+PAM’ and ‘20mer+PAM’) of each sgRNA. We used a 19-mer target sequence and added a G to our standard primer at the initial position (Supplementary Method). The ApE (A plasmid Editor) software (Davis and Jorgensen 2022, https://jorgensen.biology.utah.edu/wayned/ape/) was used to manually select multiple target sequences with higher scores (Doench et al. 2014) through an ApE-implemented ‘sgRNA Analysis’ tool.

### Plant materials and transformation

The *A. thaliana* Col-0 accession was used for the transformation. The previously generated *prk1 prk3 prk6 prk8* mutant line was used for a comparative analysis (Takeuchi and Higashiyama 2016). Plants were germinated on half-strength Murashige and Skoog (MS) solid medium at 22°C under continuous light exposure, and subsequently grown on soil under long-day conditions in a plant growth room. For transformation, each binary vector was introduced into *Agrobacterium tumefaciens* strain GV3101 through electroporation (Eporator, Eppendorf), and the T-DNA constructs were transferred via the floral dip method. For the simultaneous transformation of two constructs, two separately transformed *Agrobacterium* cultures were mixed prior to the dipping process.

For the transformation of a single construct, T1 seeds with red fluorescence were selected using a fluorescence stereomicroscope, collected in Plant Preservative Mixture, (PPM; Plant Cell Technology, Washington, D.C., USA) for surface sterilization, and sown on MS solid medium supplemented with 50 mg/L cefotaxime sodium. T1 seedlings grown on MS plates were genotyped. For the simultaneous transformation of the two constructs, T1 plants were selected on MS solid medium supplemented with 25 mg/L kanamycin, 12.5 mg/L hygromycin, and 50 mg/L cefotaxime sodium. Next, the selected plants were transferred to the soil and the recovered T1 plants were used for genotyping.

### PCR-based genotyping and direct sequencing

The primers used for PCR-based genotyping are listed in Supplementary Table S2. Genomic DNA was extracted using Nunc 96-well polypropylene DeepWell storage plates and 96-well cap mats (Thermo Fisher Scientific). Leaf samples were collected in the well containing a 3 mm tungsten carbide bead (Qiagen) and 100 µl extraction buffer (200 mM Tris-HCl, pH 8.0, 250 mM NaCl, and 25 mM EDTA), and sealed with a lid (96-well cap mats). Subsequently, the samples were crushed using a TissueLyser II (Qiagen) at 20-30 Hz for 1 min. For extracting genomic DNA, an equal volume (100 µl) of isopropanol was added to the crushed sample, followed by centrifugation at 1600 g for 20 min, and removal of the supernatant and beads by decanting. The precipitates were dried at room temperature and then dissolved in 100 µl of distilled water to obtain a purified aqueous suspension of DNA for further use in genotyping PCR. PCR amplification was performed in an 8 µl reaction mixture comprising GoTaq Green Master Mix (Promega), primers, and 1.5 µl DNA solution. Agarose gel electrophoresis analysis was performed using 5 µl of reaction mixture; 0.2-0.5 µl of the remaining PCR product was used directly for sequencing using BigDye Terminator v3.1 Cycle Sequencing Kit (Applied Biosystems) or SupreDye v3.1 Cycle Sequencing Kit (M&S TechnoSystems).

### Observation of pollen development

For the morphological analysis of pollen grains in the anther, the collected flowers were placed on a glass slide with a drop of water and squashed by pressing with a coverslip. Pollen viability staining was performed following a previously described method for the accurate phenotyping of pollen development (Takeuchi et al. 2024, Motomura et al. 2020). Epifluorescence and light microscopic analyses were conducted for squashed and stained samples, respectively, using a Ts2R microscope (Nikon).

### Analysis of double fertilization success by ovule clearing

To analyze the development of embryos and endosperms, ovules in siliques three days after pollination by wild-type pollen were cleared using chloral hydrate solution following a previously described protocol (Takeuchi et al. 2024). The samples were observed by differential interference contrast (DIC) using a Ts2R microscope (Nikon).

### Analysis of pollen tube growth and guidance

To observe pollen tubes in the pistil, mature wild-type pistils, obtained by emasculating flowers the day before, were hand-pollinated with 2-3 anthers. Pollinated pistils were fixed using ethanol mixed with acetic acid (7:3; v/v) and stained with aniline blue following a previously reported method (Maruyama et al. 2013), with slight modifications.

## Data Availability

The data underlying this study, including nucleotide and protein sequences, are available in the article and supplementary data.

## Acknowledgments

We thank Qi-Jun Chen (China Agricultural University) for plasmids, pHEE401E and pCBC-DT1T2; Tetsuya Higashiyama (The University of Tokyo) and Yoko Mizuta (Nagoya University) for discussions and supports to H.T.; Daisuke Kurihara (Nagoya University) for a technical advice on sequence design of Golden Gate assembly; Daisuke Maruyama and Daichi Susaki (Yokohama City University) for discussion on targeting *EC1* genes. This work was supported by the Program for Promoting the Enhancement of Research Universities as Young Researcher Units for the Advancement of New and Undeveloped Fields, Nagoya University.

## Author Contributions

H.T. conceived and designed this study; H.T. and S.N. generated vectors; H.T. performed genotyping PCR and established plant lines; H.T. and S.N. performed microscopy observation; H.T. wrote the manuscript with input from S.N.

