## Supplementary Method for "Development of a Versatile System to Facilitate Targeted Knockout/Elimination Using CRISPR/Cas9 for Highly Duplicated Gene Families in *Arabidopsis* Sexual Reproduction"

### Construction of CRISPR/Cas9 vectors with multiple sgRNA expression modules

For the overview of the vector construction, read the main text and Materials and Methods.

1. **Manually select specific 19-mer target sequences** using the online tool CRISPRdirect (Naito et al. 2015, http://crispr.dbcls.jp/). Another online tool, CRISPR-P (Lei et al. 2014, http://crispr.hzau.edu.cn/CRISPR2/) is also useful but need to manually check whether target sequences contain no potential off-target by seed sequence (12- or 8-mer). Note that we use 19-mer target sequences as an initial G is fixed in our standard primer design (see below).

2. **Design primers:**

For seamless cloning such as Gibson Assembly

forward primer (F primer)

*module1, 1*5*

gaaagcttggtctcgattGN(19)gttttagagctagaaatagc 58mer

*module2, 3, 4, 5, 2*5, 6*

gctagagtcgaagtagtgattGN(19)gttttagagctagaaatagc 61mer

reverse primer (R primer)

*module1, 2, 3, 4, 5, 1*5, 2*5*

cttgctatttctagctctaaaacN(19rc)Caatctcttagtcgactctac 63mer

*module6*

gtgaattcggtctcgaaacN(19rc)Caatctcttagtcgactctac 59mer

For Golden Gate assembly (GGA) cloning

forward primer (all modules)

atatgaagacggattGN(19)gttttagagctagaaatagc 55mer

reverse primer (all modules)

tattgaagacggaaacN(19rc)Caatctcttagtcgactctac 56mer

*Notes*

- *N = 19-mer target sequence without BsaI (ggtctc) and BbsI (gaagac, for GGA) sequences an initial G (or C in reverse complement (rc) primer)*
- *gttttagagctagaaatagc in the forward primer → gRNA scaffold*
  *aatctcttagtcgactctac in the reverse primer → U6-29p (reverse)*
- *ggtctc = BsaI*
- *gaagac = BbsI*

**3. PCR:**

target sequence (in F primer) + sgRNA scaffold + U6-26t + U6-29p + target sequence (in R primer)

| | |
|---|---|
| 2×KOD One (TOYOBO #KMM-201) (or favorite one) | 5.0 µl |
| $ddH_2O$ | 5.0 µl |
| template DNA (pCBC-DT1T2, x 1/200) | ~0.2 µl |
| 50 µM forward primer | ~0.2 µl (final ~1 µM) |
| 50 µM reverse primer | ~0.2 µl (final ~1 µM) |
| total | ~10 µl |

↓

98˚C, 2 min → (98˚C, 10 sec → 55˚C, 5 sec → 68˚C, 1 sec) x 20~25

↓

run on agarose gel to check amplification (~630 bp)

*Notes*

- *pCBC-DT1T2 (Xing et al. 2014, https://www.addgene.org/50590/)*
- *Negative clones with a plasmid inserted by the forward primer appear in the next step, especially when amplification is not good enough.*
- *Optional: Extraction of DNA from agarose gels or ethanol precipitation of PCR solution might reduce this background clone.*

**4. Subcloning by Gibson Assembly**

(1) digestion of pT7-module1, -2, -3, -4, -5, -6, -1*5, or 2*5 (Supplementary Table S1)
with BbsI-HF (NEB #R3539) at 37˚C
(10 µL (~1 µg) plasmid + 0.5 µL BbsI-HF in 50 µL volume)

↓

heat-inactivation by 65˚C, 20 min

(2) 0.5 µL pre-cut vector (stored at -30°C) + 0.5 µL insert (x 1/5~1/10, w/ or w/o purification)
+ 1 µL Gibson Assembly Master Mix (NEB #E2611) in PCR tube
(*minimum volume for stable handling, but colonies can be obtained even if scaled down*)

↓

50˚C, 15-30 min

↓

add ~20 µL DH5α competent cell into the PCR tube for transformation

↓

dilute with 200 µL LB and spread on LB+ampicillin plate

(3) Colony PCR (optional, but recommended, especially when used inserts w/o purification)

'U6-29p-F' primer: TTAATCCAAACTACTGCAGCCTGAC

'Hind-72F' primer: GCTGCAAGGCGATTAAG

size: ~490 bp (module1, 2, 3, 4, 5, 1*5, 2*5) / ~390 bp (module6)

(4) Sequencing by Hind-72F

Notes

- *For Golden Gate assembly to subcloning vectors, mix each pT7-moduleX vector without BbsI pre-digestion and PCR product prepared in step 3, and perform the assembly reaction.*

**5. Golden Gate assembly for integrating sgRNA modules into binary vectors**

| | |
|---|---|
| binary vector (Supplementary Table S1) (~200 ng/µl) | 0.5 µL |
| each subcloning vector (pT7-moduleX+sgRNAs) + $ddH_2O$ | 3 µL (0.5 µL each) |
| 10x T4 DNA Ligase Buffer | 0.5 µL |
| 0.1% (x10) BSA (optional) | 0.5 µL |
| T4 DNA Ligase | 0.25 µL |
| BsaI-HFv.2 (NEB #R3733) | 0.25 µL |
| total | 5 µL in PCR tube |

↓

(37˚C, 5 min → 16˚C, 5 min) x 30 → 55˚C, 10 min → 80˚C, 5 min

↓

use 1~2.5 µL for DH5α transformation and spread on LB+kanamycin plate

↓

after plasmid purification, check if proper numbers of modules are inserted by restriction enzyme digestion (*Hin*dIII + *Spe*I (or others))

Notes

- *If desired, sequencing of sgRNA modules can be performed with primers*

*module1-seq: GACATTCGATCGAGCGCAC*

*module2-seq: GACATTGAAGCCTTCGCAC*

*module3-seq: GACATCCTCATGAGTGCAC*

*module4-seq: GACATTCAAGGTTCCGCAC*

*module5-seq: GACATAGAGTTCCTCGCAC*
